# Increased glutaminolytic flux and activation of mitochondrial metabolism by BCL2 hyperactivity in lymphoma

**DOI:** 10.1101/231068

**Authors:** Kirandeep Kaur, Simar Singh, Helma Zecena, Laurent Dejean, Fabian V. Filipp

## Abstract

B-cell lymphoma 2 (BCL2) is an important apoptosis regulator during developmental and pathological states, and its overexpression is a key feature of several malignancies. Genomic data from The Cancer Genome Atlas (TCGA) reveals significant somatic copy number amplification, overexpression, and/or elevated protein activity of BCL2 in 50 % of diffuse large B-cell lymphoma (DLBC) patients. While its canonical role in mitochondria-directed apoptosis is well established, the effect of BCL2 on transcriptional and metabolic networks remains elusive. Using an established lymphocytic pro-B-cell line overexpressing BCL2, we identified dysregulated transcriptional and metabolic networks by transcriptomic profiling arrays. Elevated BCL2 levels affect transcription factor complexes and mitogenic programs of NF-κB/REL, HIF1A/ARNT, AP1, E2F, and STAT factors. Using stable isotope-assisted metabolic flux measurements we quantify that elevated BCL2 expression increases carbon utilization boosting cellular proliferation. Tumorigenic overexpression of BCL2 significantly increases glycolytic flux, glutaminolysis, and anaplerotic flux into the TCA cycle. At the same time, the mitochondrial acetyl-CoA pool is separated from the glycolytic one by inactivating the pyruvate dehydrogenase complex via transcriptional regulation of pyruvate dehydrogenase kinase (PDK3). As compensatory fuel, mitochondrial TCA cycle metabolism is supported by asparagine synthase (ASNS) and oxidative glutaminolysis creating targets for small molecule inhibition of glutaminase. Lymphoma cells overexpressing BCL2 contained more mitochondrial mass and were more sensitive to L-glutamine deprivation and glutaminase inhibition. Cells overexpressing a mutant BCL2 G145E, which is incapable of binding BH domain members, failed to increase proliferation, glycolysis, or glutaminolysis. Taken together, the oncogene BCL2 has the ability to ramp up a metabolic phenotype supporting proliferation independent of its anti-apoptotic role. The cellular model of BCL2 activation supports NF-κB-positive subtypes of DLBC and identifies metabolic bottlenecks with dependency on anaplerotic flux as an actionable BCL2 effector network in cancer.

## Introduction

The B-cell lymphoma 2 (BCL2, Gene ID: 596) family of proteins are essential regulators of the intrinsic, mitochondrial pathway of apoptosis ^1^. They are characterized by the presence of BCL2 homology (BH) domains which interact with one another to regulate essential cellular processes such as survival pathways, apoptotic initiation, cell cycle propagation, mitochondrial activity, or autophagy transitions ^2^. Phylogenetically, members of the BCL2 family classifies as either pro- or anti-apoptotic BCL2 homologs, members with canonical BH domains, or those with non-canonical BH domains. Of the pro-apoptotic BH member the BCL2 associated X apoptosis regulator (BAX, Gene ID: 581) and BCL2 antagonist/killer 1 (BAK, Gene ID: 578) are interaction partners and direct effectors of mitochondrial outer membrane permeabilization (MOMP). In contrast, in case there is no interaction with pro-apoptotic BH members, BCL2 and BCL2 like 1 (BCL2L1, better known as BCL-XL, Gene ID: 598) exert their anti-apoptotic effect through inhibitory interactions with pro-apoptotic members such as BAX.

BCL2 was first described as part of the t(14:18) chromosomal translocation event in B-cell follicular lymphoma ^3^, where it functioned to block apoptosis^4, 5^.The translocation of BCL2 to the Ig heavy chain locus in chromosome 14 drives its constitutive expression ^6-8^. Subsequent studies demonstrated that BCL2 is overexpressed in several hematological cancers ^9^ and may support cancer cell survival through chemotherapeutic resistance ^10^ and regulation of autophagy^11, 12^. Clinically, trials of BCL2 inhibitors such as ABT-199 (venetoclax, PubChem CID: 49846579) have shown promise in the treatment of hematological cancers and immune diseases ^13-16^.

Cellular proliferation, including both physiologic lymphoid cell expansion and pathologic malignancy, requires a metabolic phenotype that supports macromolecular biosynthesis. Mitochondrial activity, electron redox transfer in the inner mitochondrial membrane, and oxidative phosphorylation play key roles in satisfying the energy demands of growing and diving cells. Despite a growing understanding of the role of BCL2 in mitochondria-directed apoptosis, little is known about whether BCL2 regulates metabolism under non-apoptotic conditions.

Interleukin 3 (IL3, Gene ID: 3562) dependent FL5.12 pro-B cells derived from murine fetal liver ^17^ have served as a useful model for studying B-lymphocyte development ^18^, oncogenesis^19, 20^, apoptosis ^21, 22^, and BCL2 binding events^23, 24^. Overexpression of BCL2 in these cell lines ^25^ leads to increased survival following IL3 deprivation, and increased tumorigenesis *in vivo* ^20^. Using the FL5.12 cell line, and stably expressing BCL2^20, 26^ or a mutant of BCL2 unable to bind BH members^23, 27-30^, we sought to quantify the transcriptional and metabolic phenotype of BCL2 overexpression using metabolic flux and proliferation assays.

## Methods

### Cell Culture

Lymphocytic FL5.12 pro-B-cell lines including a parental lymphocytic cell line with BCL2 wild type ^17^, a tumorigenic lymphoma cell line with BCL2 overexpression ^25^, and a BCL2(G145E) mutant with loss of binding of BH domain interaction partners ^23^ were cultured in minimal essential media (MEM, 15-010-CV, Corning, Corning, NY) supplemented with 10.0 % fetal bovine serum (FBS, 35-010-CV, Corning, Corning, NY), 1.0 % penicillin-streptomycin solution (PS, 30-002-CI, Corning, Corning, NY), 1.0 % MEM vitamins (MEM VIT, 25-020-CI, Corning, Corning, NY), 0.8 ng/mL of interleukin 3 (IL3, I4144, Sigma Merck, Darmstadt, Germany), 2.0 μL of 2-mercaptoethanol (M6250-100ML, Sigma Merck, Darmstadt, Germany), 1.0 g/L D-glucose (GLC, G7021-1KG, Sigma Merck, Darmstadt, Germany), and 2.0 mM L-glutamine (GLN, 25-005-CI, Corning, Corning, NY) at 37 °C (310 K) with 5.0 % carbon dioxide (CO2, CD50, Praxair, Danbury, CT).

### Cell size and proliferation

Cell diameter and proliferation rates of normal and cancer cells were quantified by automated imaged-based cytometry. Cells in suspension were harvested and mixed 1:1 with a 0.4 % solution (w/v) trypan blue (25-900-CI, Corning, Corning, NY), and pipetted into disposable counting chambers (1450003, Bio-Rad, Hercules, CA) for counting and image analysis. Cell diameter and proliferation rates measurements of live cells in exponential growth were obtained in an automated tissue cell counter (TC20, 145-0102, Bio-Rad, Hercules, CA). Multi-planar bright-field digital images were automatically collected, quantified, and assessed for cell number and diameter. Cell proliferation rates were calculated and densities validated from the live cells per flask (N=6) over a 3-day time course of a proliferation assay experiment. Proliferation data based on different initial seeding densities was LOG2 transformed to fit a linear regression model with an explained variation R^2^ above 0.970.

### Flow cytometry and mitochondrial stains

Cells were analyzed for mitochondrial content, membrane potential, and matrix oxidant burden as previously described ^31^. Cells were collected by centrifugation, counted, and aliquots corresponding to 500,000 cells were incubated protected from light with 200 nM MitoTracker green fluorescent mitochondrial stain (MTG, M7514, Life Technologies, Thermo Fisher Scientific, Carlsbad, CA), 13.3 nM tetramethylrhodamine ethyl ester (TMRE, T669, Life Technologies, Thermo Fisher Scientific, Carlsbad, CA), or 6.6 μM MitoSOX red mitochondrial superoxide indicator (MitoSOX, M36008, Life Technologies, Thermo Fisher Scientific, Carlsbad, CA). Cells were collected, centrifuged, and resuspended in phosphate buffered saline (PBS, 46-013-CM, Corning, Corning, NY) supplemented with 2.0 % FBS twice before being analyzed on an LSR II flow cytometer (BD Biosciences, San Jose, CA) at a flow rate at least 500 events per second. 100,000 events per sample were recorded and samples were analyzed in triplicate (N=3) for each data point. FlowJo (V10, FlowJo, Ashland, OR) was used for data analysis.

### Custom-designed gene expression profiling arrays for high-throughput RT-QPCR

We generated custom-designed profiling arrays to validate differential expression of metabolic genes and transcription factors in the *in vitro* lymphoma progression model of BCL2 activation. Total RNA was extracted from FL5.12 murine pro-lymphocyte B-cells using NucleoSpin RNA Plus Columns (740984.25, Macherey-Nagel, Düren, Germany). At least three biological replicates of RNA samples were analyzed per condition. The concentration of RNA was determined using a micro-volume plate (Take3, BioTek, Winooski, VT) and a multi-mode microplate reader (Synergy HT, BioTek, Winooski, VT) multi-mode microplate reader (Synergy HT, BioTek, Winooski, VT). One microgram of RNA was used to synthesize complementary DNA (cDNA) using qScript cDNA SuperMix (95048-500, Quanta Biosciences, Beverly, MA). cDNA corresponding to 1/5 of the first strand synthesis was mixed with 500 nM custom-designed forward and reverse primers (Sigma Genosys, The Woodlands, TX) and PerfeCTa SYBR Green FastMix (95072-05k, Quanta Biosciences, Beverly, MA) and analyzed on a high-throughput real-time (RT) quantitative polymerase chain reaction (QPCR) System (ECO, EC-101-1001R, Illumina, San Diego, CA). Custom-designed gene expression profiling arrays were analyzed using the ΔΔCT method. RT-QPCR threshold cycle (CT) values were normalized using three different housekeeping genes (HKG), ribosomal protein S13 (RPS13, Gene ID: 6207), TATA box binding protein (TBP, Gene ID: 6908), and polymerase (RNA) II (DNA directed) polypeptide A (POLR2A, Gene ID: 5430). The difference threshold cycle value (ΔCT) of any gene of interest (GOI) to the average housekeeping value was calculated using the formula ΔCT(GOI) = CT(GOI) — AVERAGE(CT(HKG)) for each cell line. In addition, change in expression of any gene of interest was monitored by calculating ΔΔCT(GOI) = ΔCT(GOI^CONDITION^) — ΔCT(GOI^WT^) with CONDITION being cell lines overexpressing BCL2 or BCL2(G145E) and WT the reference lymphocytic pro-B-cell line. RT2 gene array profiles were normalized, separated according to differential expression between conditions in univariate T-tests with a random variance model using a p-value cut-off below 5.00E-02, and ranked with LOG2 fold-change between specimens considered significant.

### Somatic copy number alteration analysis

The tool GISTIC, genomic identification of significant targets in cancer, 2.0.21 ^32-34^ was used to identify genomic regions that are significantly gained or lost across a set of 48 paired normal and tumors samples of TCGA DLBC data set. We executed GISTIC 2.0.21 on Agilent SNP 6.0 gene expression microarrays G4502A_07_01, UNC Chapel Hill, NC. GISTIC 2.0.21 distinguishes arm-level events from focal events at a broad length cutoff of 0.7. Events whose length was greater or less than 50% of the chromosome arm on which they resided were called arm-level or focal events, respectively, and these groups of events were analyzed separately. The data was concordant to segmented level 3 data publically available at the TCGA data portal. Since GISTIC 2.0.21 uses ratios of segmented tumor copy number data relative to normal samples as input, segmented level 3 data aligned to HG19 served as input for analysis runs. For significant loci and genes a cutoff p-value of 0.05 and q-value of 0.1 was applied, and concordance determined by overlaying whole-genome sequencing and SNP data. All experiments on SCNAs were carried out at a confidence level of 0.99.

### Cell culture for metabolomics flux measurements

For metabolite quantification and stable isotope tracing, 100,000 cells were seeded in tissue culture flasks with vented caps (430639, Corning, Corning, NY) in N=6 replicate. For stable isotope D-glucose and L-glutamine labeling experiments, MEM media was supplemented with 1 g/L [U-^13^C_6_] D-glucose ([U-^13^C_6_] GLC, 389374-2G, Sigma Isotec, Miamisburg, OH) or 2 mM [U-^13^C_5_] L-glutamine ([U-^13^C_5_] GLN, 605166-500MG, Sigma Isotec, Miamisburg, OH). After 24 h, cell suspensions were transfer to microcentrifuge tubes (MT-0200-BC, Biotix, San Diego, CA) and centrifuged for 5 min at 4 °C (277 K) and 300 g in a refrigerated centrifuge (X1R Legend, Sorvall, Thermo Fisher Scientific, Waltham, MA) using a fixed-angle rotor (F21-48x1.5, Sorvall, Thermo Fisher Scientific, Waltham, MA). For exometabolome analysis, 40 μL of supernatant containing condition media was transferred to microcentrifuge tubes and dried by vacuum centrifugation in a speedvac concentrator (DNA120OP115, Savant, Thermo Fisher Scientific, Waltham, MA) overnight. The remaining supernatant was aspirated and the cell pellets frozen in liquid nitrogen before storage at -80 °C (193 K).

### Metabolite extraction

Frozen cell pellets were thawed on ice for 10 min before addition of 1 mL -20 °C (253 K) cold extraction solvent containing acetonitrile/isopropanol/water (3:3:2) acetonitrile (ACN, 34998-4L, Sigma Merck, Darmstadt, Germany), isopropanol (IPA, 34965-1L, Sigma Merck, Darmstadt, Germany), water (H2O, 46-000-CI, Corning, Corning, NY). Samples were then vortexed 5 times for 15 s and frozen on dry ice for 20 min and the freeze/thaw/vortex cycle repeated twice. Extracted cell suspension or media supernatants were dried via vacuum centrifugal evaporation and stored at -80 °C (193 K) before analysis.

### Metabolite derivatization

Dried, extracted cell pellets or media supernatants were derivatized by addition of 20 μL of 2.0 % methoxyamine-hydrochloride in pyridine (MOX, TS-45950, Thermo Fisher Scientific, Waltham, MA) followed by 90min incubation in a digital heating shaking drybath (8888-0027, Thermo Fisher Scientific, Waltham, MA) at 30 °C (303 K) and 1100 rpm. 90 μL of N-methyl-N-(trimethylsilyltrifluoroacetamide (MSTFA, 394866-10X1ML, Sigma Merck, Darmstadt, Germany) was added and samples incubated at 37 °C (310 K) and 1100 rpm for 30 min before centrifugation for 5 min at 14,000 rpm and 4C. The supernatant was transferred to an autosampler vial (C4000LV3W, Thermo Fisher Scientific, Waltham, MA) with caps (C5000-53B, Thermo Fisher Scientific, Waltham, MA) for separation by gas chromatography (GC, TRACE 1310, Thermo Fisher Scientific, San Jose, CA) coupled to a triple-quadrupole GC mass spectrometry system (QQQ GCMS, TSQ8000EI, TSQ8140403, Thermo Fisher Scientific, San Jose, CA) for analysis.

### GCMS for metabolomics

Samples were analyzed on a QQQ GCMS system equipped with a 0.25 mm inner diameter, 0.25 μm film thickness, 30 m length, low polarity phase, 5% diphenyl/95% dimethyl polysiloxane capillary column (TraceGOLD TG-5MS, 26098-1420, Thermo Fisher Scientific, Waltham, MA) and run under electron ionization at 70 eV. The GC was programed with an injection temperature of 250 °C (523 K) and splitless injection volume of 1.0 μl. The GC oven temperature program started at 50 °C (323 K) for 1 min, rising to 300 °C (573 K) at 10 K/min with a final hold at this temperature for 6 min. The GC flow rate with helium carrier gas (HE, HE 5.0UHP, Praxair, Danbury, CT) was 1.2 mL/min. The transfer line temperature was set at 290 °C (563 K) and ion source temperature at 295 °C (568 K). A range of 50-600 mass over charge (m/z) was scanned with a scan time of 0.25 s.

### Metabolomics data processing

Metabolites were identified using metabolite retention times and fragmentation patterns in TraceFinder (v3.3, Thermo Fisher Scientific, Waltham, MA). Identified metabolites were quantified using the selected ion count peak area for specific mass ions, and standard curves generated from reference standards run in parallel. The mean, standard deviation, and 95.0 % confidence interval for each quantified metabolite was calculated for each cell line and treatment condition. A two-sample homoscedastic student’s t-test was used to compare treatment conditions of each metabolite and each cell line.

### ^13^C stable isotope tracing and metabolic flux quantification

To quantify metabolic fluxes from substrates into metabolites, the mass isotope distribution vector (MDV) for known fragments of carbon backbone labeled amino acids and carboxylic acids was retrieved and ^13^C tracer-to-tracee ratios were calculated ^35-37^. GCMS MDV data from fractionally labeled [U-^13^C] D-glucose or -L-glutamine samples was quantified to determine pool and isotopomer distribution of intracellular metabolites. Identified fragments contained either the whole carbon skeleton of the metabolite or resulted from a loss of the carboxyl carbon, or for some amino acids contained only the backbone minus the side-chain. For each fragment, mass ion counts were retrieved for the lightest isotopomer (M+0, without any heavy isotopes), and cumulative isotopomers for heavier mass counts with increasing mass units (M+1 up to M+6) relative to M0. Mass ion counts were normalized by dividing by the sum of M0 to M6, and corrected for the natural abundance of heavy isotopes of the elements H, N, O, Si, and C. By using probabilistic matrix-based multiplication one arrives at normalized MDVs corrected for naturally occurring isotopes in atoms of fragments of the metabolite backbone. ^13^C-labeling data is expressed as fraction of the MDV and corresponds to stable isotope enrichment per carbon in a measured metabolite for a set of biological replicates with number of experiments N=6 for each condition. ^13^C-labeling data is converted into metabolic flux from carbon source by dividing by percent labeling of respective carbon source, 50 % [U-^13^C_6_] of total 2.0 g/L D-glucose or 100 % [U-^13^C_5_] of total 2.0 mM L-glutamine. A two-sample student’s t-test with a minimum significance level α=5.00E-02 was used to compare average stable isotope enrichment per carbon of each metabolite between conditions.

### L-glutamine deprivation and glutaminase inhibition

Cells were plated in white 96-well assay plates at a density of 10,000 cells/well in 100 μL of media containing either no L-glutamine, 2.0 mM L-glutamine, or 2.0 mM L-glutamine with 1.0 mM selective glutaminase inhibitor bis-2-(5-phenylacetamido-1,3,4-thiadiazol-2-yl)ethyl sulfide (BPTES, SML0601, Sigma Merck, Darmstadt, Germany). After 24 h, cells were equilibrated at room temperature before being treated with 100 μL/well of CellTiter-Glo luminescent cell viability assay (CTG, G7571, Promega, Madison, WI). Cells were then shaken for 2.0 min and incubated for 8 min protected from light before being analyzed in a multi-mode microplate reader (Synergy HT, BioTek, Winooski, VT).

## Results

### Somatic activation of BCL2 in lymphoma

In lymphoma, one of the most frequent somatic copy number alteration (SCNA) events is arm-level amplification of chromosome 18 with a recurrence frequency of 30 % in The Cancer Genome Atlas (TCGA) dataset (Figure 1A). In DLBC, significant arm-level SCNAs include: chr18pq, 30 %, 7.66E-04; chr7pq, 26 %, 4.57E-04; chr3pq, 23 %, 1.43E-02; chr11pq, 19 %, 1.43E-01; chr21q, 21 %, 1.08E-01; chr12q, 19 %, 2.83E-02; chr8q, 18 %, 1.75E-01; and chr1q, 13 %, 2.48E-01. These SCNA events enhance function of oncogenes and tumor suppressors in lymphoma (Figure 1A-B). Specifically, a sharply-defined focal region around chromosome band 18q21 (genome coordinates chr18:48582939-78077248) is significantly amplified with a q-value of 0.065049. Further, multi-omics integration reveals that BCL2 activation is often observed at multiple different levels including copy number, gene expression, and/or protein expression (Figure 1C). Elevated BCL2 activity has a negative effect on overall decreased patient survival (Figure 1C-D) ^38-42^.

**Figure 1:**
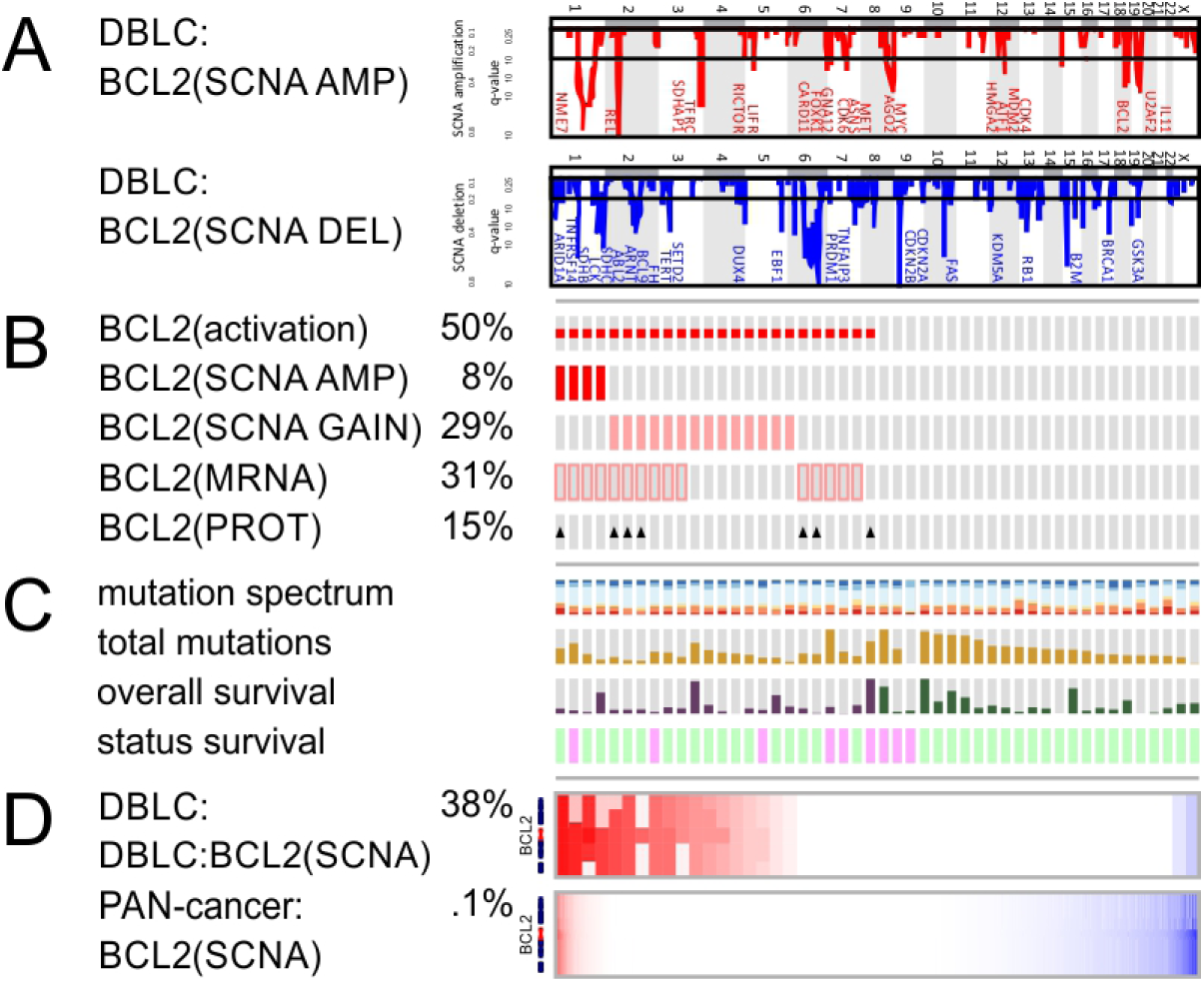
Frequent somatic copy number aberration of chromosome 18 in B-cell lymphoma results in amplification and somatic activation of BCL2. A) Landscape of somatic copy number aberrations (SCNAs) of diffuse large B-cell lymphoma (DLBC) with significant amplifications (red) and deletions (blue). B) Frequency and mechanism of somatic activation of B-cell lymphoma 2 (BCL2, Gene ID: 596) in DLBC. C) Mutation spectrum and impact of BCL2 activation on overall survival. D) Comparison of somatic aberrations of BCL2 in DLBC vs all cancer tissues (PAN-cancer) cohort in The Cancer Genome Altas (TCGA).

Somatic BCL2 amplification shows significant co-occurrence with other oncogenic drivers observed in DLBC and highlights synergistic effects of structural somatic events in disease initiation and progression. Together with somatic amplification of BCL2 the following SCNA events are observed and constitute some of the most frequent and significant focal SCNA events in DLBC: deletion of cyclin dependent kinase inhibitor 2A and 2B (CDKN2A, Gene ID: 1029, CDKN2B, Gene ID: 1030, chr9p21); amplification of proto-oncogene and NF-κB interaction partner REL (REL, REL proto-oncogene, NF-κB subunit, Gene ID: 5966, chr2p16), B-cell CLL/lymphoma 11A (BCL11A, Gene ID: 53335, chr2p16), B-cell CLL/lymphoma 6 (BCL6, Gene ID: 604, chr3q27), non-receptor tyrosine kinase ABL proto-oncogene 2 (ABL2, Gene ID: 27, chr1q25) (Figure 1, Supplementary table 1). Taken together, genomic concertation and correlation of focal chromosome aberrations at the copy number level with focal BCL2 amplification emphasizes tissue-specific oncogenic driver pathways in lymphoma.

B-cell activation, selection, and maturation rely on apoptotic and pro-survival pathways mediated by BH domain proteins. BCL2 holds a tissue-specific gene expression program in the B-cell lineage and in lymphatic cancers. We therefore, sought to compare incidence of BCL2 activation in lymphomas with all other cancer tissues. Somatic activation of BCL2 is observed in less than 0.1 % of 10944 specimens across 37 pan-cancer tissues. There are two cases of gene fusions of BCL2 with a chromatin remodeler and a neural regulator: ATRX-BCL2 in lower grade glioma and NEDD4L-BCL2 fusion pair in breast cancer in combination with SCNAs activate the gene product. Therefore, across all cancers, DLBC stands out as cancer tissue with frequent arm-level amplification of chromosome 18 and distinct transcriptional activation of BCL2. In comparison to all other cancers, DLBC with a cohort size of 48 specimens is highly enriched for somatic amplification of chromosome 18. The detected DLBC cases represent about half of all highly amplified focal amplification cases observed across more than 10,000 cancer specimens.

### Enrichment of inflammatory and metabolic pathways in specimens with BCL2 activation

In order to prioritize pathways affected by BCL2 activation, the DLBC patients were divided into two sub-cohorts: on the one hand, specimens were assigned to an unaltered sub-cohort with normal BCL2 status. On the other hand, a sub-cohort was defined with BCL2 activation based on somatic copy number amplification and differential expression of RNA-Seq V2 RSEM data. Using these two sub-cohorts, we queried whether activation of BCL2 resulted in enrichment of significantly deregulated transcripts and proteins. In protein arrays, BCL2 overexpression was maintained at the protein level with a p-value of 3.77E-07 (Figure 2A). This is an important data point to validate, since the query cohort was selected exclusively based on genomic and transcriptional activation. One of the strongest activated proteins was the L-glutamine-hydrolyzing metabolic enzyme asparagine synthetase (ASNS, Gene ID: 440) (Figure 2B). Remarkably, focal SCNA amplification on chr7q21 of ASNS (Figure 1A) occurs in 29.2 *%* of patients and coincides with SCNA of BCL2 with a p-value of 1.70E-02. In addition, a set of cycle genes was enriched including cyclin dependent kinase inhibitor 1B (CDKN1B, Gene ID: 1027), proliferating cell nuclear antigen (PCNA, Gene ID: 5111), cyclin B1 (CCNB1, Gene ID: 891), and cyclin E1 (CCNE1, Gene ID: 898) with a p-values below 1.00E-02 (Figure 2C-D). Cyclin dependent kinase inhibitor protein CDKN1B binds to and prevents the activation of cyclin E-CDK2 or cyclin D-CDK4 complexes. In the altered, BCL2-activated gene set, there was significant enrichment of metabolic pathways including amino acid processing, oxidative phosphorylation, hypoxic and glycolytic metabolism with p-values below 1.0E-10 (Table 1). Significantly enriched transcriptional networks included complexes of signal transducer and activator of transcription (STATs, specifically 3/5A/5B), nuclear factor kappa B (NF-κB), NF-κB subunit REL proto-oncogene (REL), JUN proto-oncogene/FOS proto-oncogene complex (AP1), yin yang (YY) transcription factor, hypoxia inducible factor (HIF), and aryl hydrocarbon receptor nuclear translocator (ARNT) transcription factor families with p-values below 2.0E-02. In addition, pathways in cancer, interferon response, inflammatory response, janus kinase (JAK)-STAT signaling, and the tumor protein p53 pathway (TP53, Gene ID: 7157) were enriched with p-values below 1.4E-03. The analysis indicates that BCL2 has the ability to activate a specific effector network impacting cell cycle, inflammation, and metabolism.

**Figure 2:**
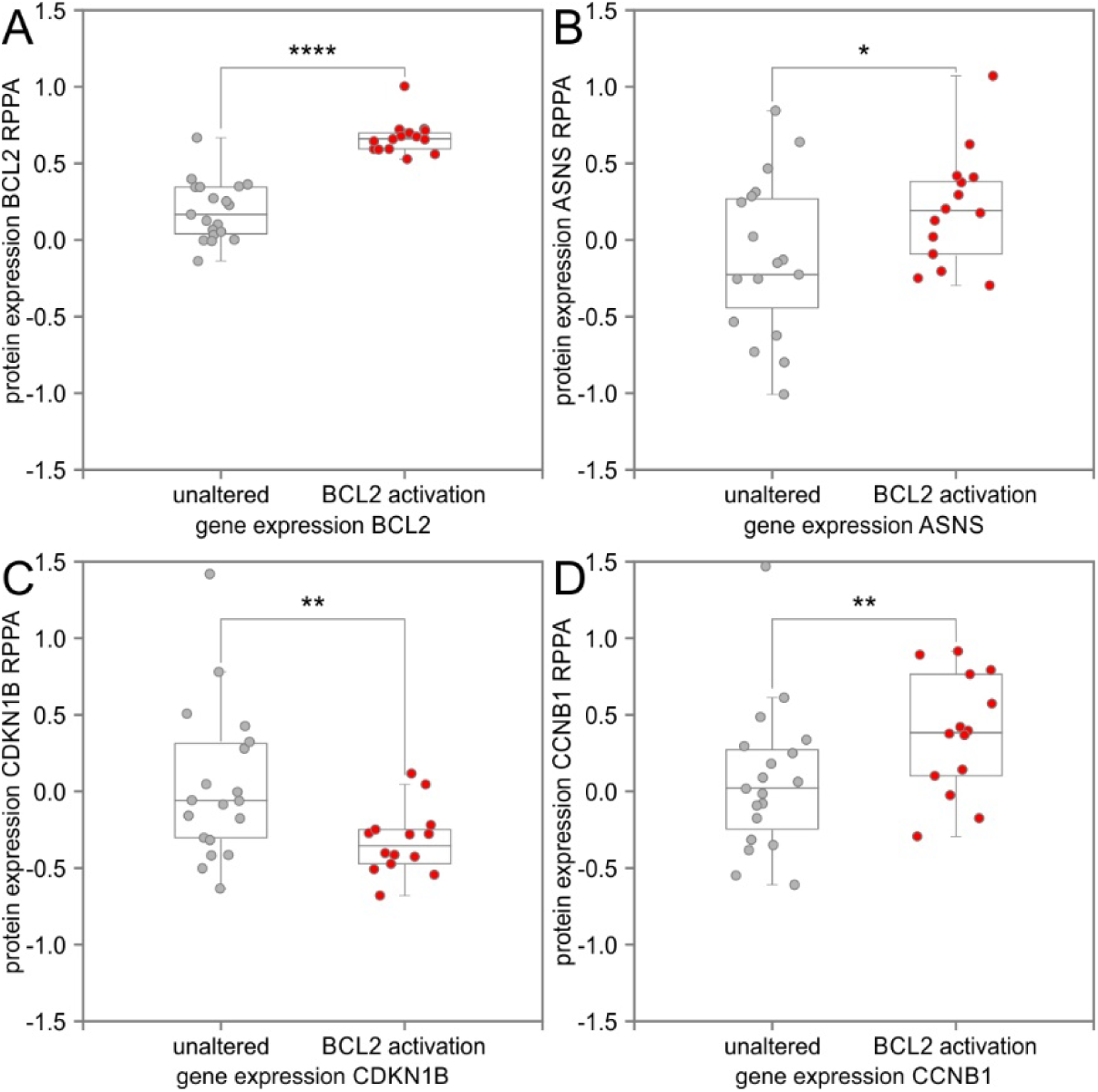
BCL2 activation significantly modulates metabolic and cell cycle regulators at the protein level. Protein expression level of lymphoma patients with unaltered BCL2 status was compared to patients with BCL2 activation. A) Specimens with BCL2 activation were selected based on altered somatic copy number or differential expression and had significantly higher BCL2 protein levels. B) asparagine synthetase (ASNS, Gene ID: 440) protein levels were elevated in lymphoma patients with genomic or transcriptomic BCL2 activation. C) The cyclin dependent kinase inhibitor 1B (CDKN1B, Gene ID: 1027) controls cell cycle progression and had lower median levels in patients with BCL2 activation. Degradation or loss of this protein is required for transition to a highly proliferative state. D) Cell cycle regulator cyclin B1 (CCNB1, Gene ID: 891) showed higher median protein levels with BCL2 activation. Analysis of protein expression levels was performed on reverse phase protein arrays (RPPA) of The Cancer Proteome Atlas (TCPA). Quartile box plots illustrate distribution following data normalization. Significance level of differential expression is indicated by asterisks according to p-value thresholds (* p-value < 5.00E-02, ** p-value < 1.00E-02, **** p-value < 1.00E-04).

**Table 1:** Gene ontology and pathways enriched with BCL2 activation

| NAME | FRAMEWORK | SIZE | NES | p-value | q-value |
| --- | --- | --- | --- | --- | --- |
| Regulation of amino acid process | GO | 56 | 2.41 | 0.0E+00 | 1.2E-03 |
| TCA cycle and respiratory chain | GO | 114 | 2.12 | 0.0E+00 | 3.1E-03 |
| Amine metabolic process | GO | 85 | 1.82 | 0.0E+00 | 5.6E-02 |
| Metabolism of amino acids | GO | 136 | 1.69 | 0.0E+00 | 7.0E-02 |
| Precursor metabolites and energy | GO | 238 | 1.71 | 0.0E+00 | 8.8E-02 |
| Cellular amide metabolic process | GO | 594 | 1.46 | 1.2E-03 | 2.0E-01 |
| Oxidative phosphorylation | KEGG | 196 | 2.72 | 0.0E+00 | 0.0E+00 |
| Epithelial mesenchymal transition | KEGG | 173 | 2.69 | 0.0E+00 | 0.0E+00 |
| Coagulation | KEGG | 92 | 2.64 | 0.0E+00 | 0.0E+00 |
| Interferon gamma response | KEGG | 189 | 2.57 | 0.0E+00 | 0.0E+00 |
| Interferon alpha response | KEGG | 94 | 2.57 | 0.0E+00 | 0.0E+00 |
| Complement | KEGG | 163 | 2.52 | 0.0E+00 | 0.0E+00 |
| Inflammatory response | KEGG | 160 | 2.47 | 0.0E+00 | 0.0E+00 |
| TNFα signaling via NFκB | KEGG | 181 | 2.28 | 0.0E+00 | 0.0E+00 |
| Allograft rejection | KEGG | 175 | 2.15 | 0.0E+00 | 0.0E+00 |
| IL2 STAT5 signaling | KEGG | 183 | 2.02 | 0.0E+00 | 3.4E-04 |
| Hypoxia | KEGG | 166 | 1.90 | 0.0E+00 | 7.7E-04 |
| Glycolysis | KEGG | 170 | 1.91 | 0.0E+00 | 8.3E-04 |
| Apoptosis | KEGG | 148 | 1.86 | 0.0E+00 | 9.6E-04 |
| TP53 pathway | KEGG | 180 | 1.67 | 1.3E-03 | 5.9E-03 |
| IL6 JAK STAT3 signaling | KEGG | 73 | 1.63 | 4.6E-03 | 9.2E-03 |
| Fatty acid metabolism | KEGG | 132 | 1.55 | 7.2E-03 | 1.8E-02 |
| Angiogenesis | KEGG | 29 | 1.58 | 1.8E-02 | 1.4E-02 |
| \$STAT5A | TF | 174 | 1.73 | 0.0E+00 | 1.3E-02 |
| \$NFKB1 | TF | 195 | 1.70 | 0.0E+00 | 1.4E-02 |
| \$STAT5B | TF | 148 | 1.67 | 0.0E+00 | 1.5E-02 |
| \$E2F | TF | 200 | 1.66 | 0.0E+00 | 1.6E-02 |
| \$AP1 | TF | 170 | 1.64 | 1.4E-03 | 1.6E-02 |
| \$HIF1A | TF | 225 | 1.63 | 1.6E-03 | 6.6E-03 |
| \$ARNT | TF | 257 | 1.62 | 5.0E-03 | 1.5E-02 |
| \$YY1 | TF | 206 | 1.61 | 2.7E-02 | 1.9E-02 |

### Validation of dysregulated metabolic and transcriptional networks by transcriptomic profiling arrays

We utilized custom-designed gene expression profiling arrays based on identified genomic and transcriptomic alterations with BCL2-activation in DLBC. Differential gene expression analysis of the *in vitro* lymphoma progression model of BCL2 activation validated important regulatory molecules and control points of the metabolic and transcriptional machinery related to the oncogenic effector network of BCL2. We identified 28 transcripts of 75 tested target genes to be significant differentially expressed in response to BCL2 overexpression with p-values below 1.0E-05 (Figure 3, Supplementary table 2). The differential gene expression analysis took into consideration: basal expression level and directionality of regulation highlighting transcription factors (Figure 4A) and functionally redundant metabolic isoenzymes (Figure 4B). For enhanced clarity, two-dimensional plotting of gene expression values of multiple conditions facilitates visual inspection of differential regulation (Figure 3), while bar graphs in direct comparison of conditions emphasize basal expression level and trends in directionality (staggered presentation of gene expression of pro-B-cell line overexpressing BCL2, pro-B control cell line, and BCL2(G145E) mutant) (Figure 4). For instance, the target gene ASNS has higher gene expression in the pro-B-cell line overexpressing BCL2 but lower expression in the BCL2(G145E) mutant. Its gene expression values are therefore plotted above and below the diagonal, respectively, against the pro-B control cell line (Figure 3A). The data indicates that ASNS expression is favored upon BCL2 overexpression but decreased if BH domain interactions are lost in the mutant. Many metabolic enzymes showed opposite directionality of regulation following BCL2 or BCL2(E145G) mutant overexpression including asparagine synthetase glutamine-hydrolyzing (ASNS, Gene ID: 440), hexokinase 2 (HK2, Gene ID: 3099), glucose-6-phosphate dehydrogenase (G6PD, Gene ID: 2539), transketolase (TKT, Gene ID: 7086), glyceraldehyde-3-phosphate dehydrogenase (GAPDH, Gene ID: 2597), glyceraldehyde-3-phosphate dehydrogenase, spermatogenic (GAPDHS, Gene ID: 26330), lactate dehydrogenase A (LDHA, Gene ID: 3939), and pyruvate dehydrogenase kinase 3 (PDK3, Gene ID: 5165) (Figure 3A, Figure 4B). HIF/ARNT and STAT transcription factor recognition sites are common to this set of metabolic target genes. Regulators of the L-glutamine metabolite pool, ASNS, and pyruvate flux, PDK3, take pivotal roles standing out as strongest, most significantly regulated target genes (Figure 3A). In contrast, GCK, HK3, PFKM, PFKP, PDHA1, PDHB, and PDK1 were down-regulated following BCL2 overexpression (Figure 3B). Further, prominent transcription factors, RELA proto-oncogene (RELA, NF-κB subunit, NF-κB3, p65, Gene ID: 5970), nuclear factor kappa B subunit 1 (NFKB1, p105, p50, Gene ID: 4790), hypoxia inducible factor 1 alpha subunit (HIF1A, BHLHE78, Gene ID: 3091, aryl hydrocarbon receptor nuclear translocator (ARNT, Gene ID: 405, signal transducer and activator of transcription 5A (STAT5A, Gene ID: 6776), signal transducer and activator of transcription 5B (STAT5B, Gene ID: 6777), signal transducer and activator of transcription 6 (STAT6, Gene ID: 6778), and Jun proto-oncogene, AP1 transcription factor subunit (JUN, AP1, Gene ID: 3725), showed significant regulation in both test conditions (Figure 3A-C, Figure 4A). The *in vitro* analysis validated activation of heterodimeric transcription factor complex components NFKB1/REL, HIF1A/ARNT, AP1, E2F, and STAT5A/5B/6 with BCL2 activation in agreement with the genomic data (Table 1, Figure 4A). Gene ontology and pathway analysis pointed toward activation of amino acid metabolism, enhanced glycolysis, and TCA cycle with BCL2 activity. The profiling of metabolic enzymes validated this finding at the transcriptional level and showed up-regulation of key metabolic mediators of glycolysis, TCA cycle, and anaplerosis (Table 1, Figure 2B, Figure 4B).

**Figure 3:**
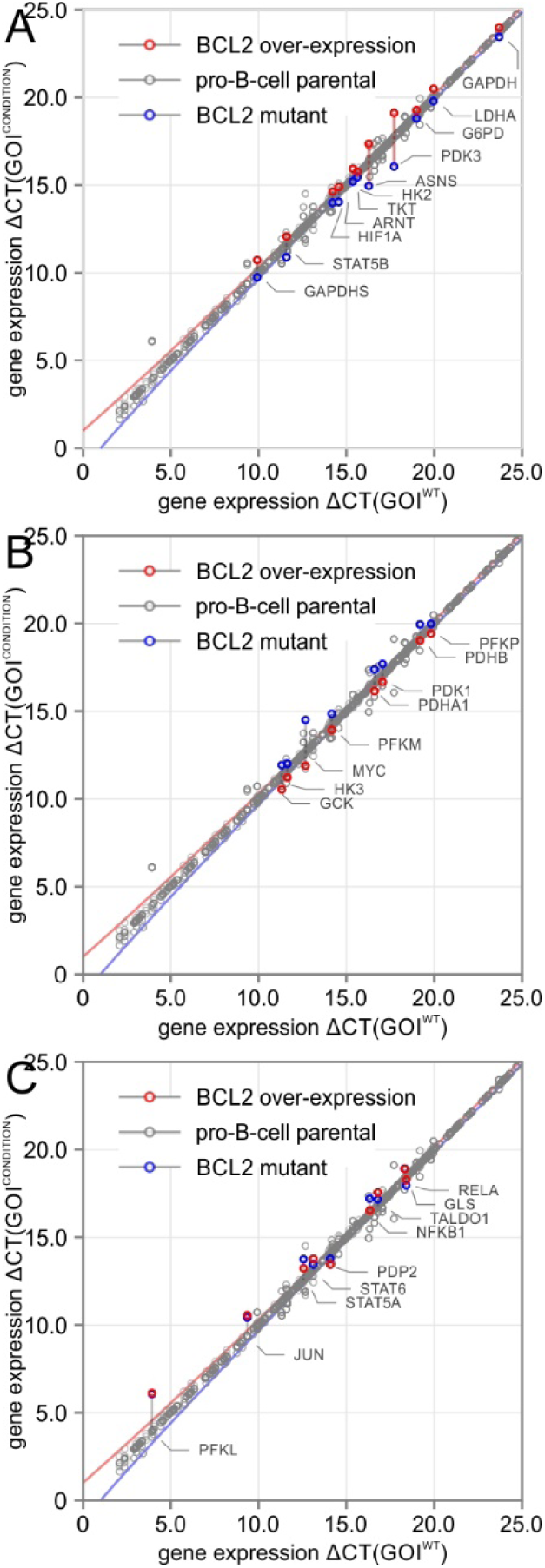
Elevated BCL2 expression results in separation of the mitochondrial acetyl-CoA pool from the glycolytic one by transcriptional regulation of the pyruvate dehydrogenase complex via PDK3. Real-time quantitative polymerase chain reaction (RT-QPCR) profiling arrays of central carbon metabolic enzymes and their regulators validate significant deregulation of mitochondrial metabolism at the gene expression level. Gene expression level of the gene of interest (GOI) is shown as negative threshold cycle (-ΔCT) relative to the house keeping gene for the parental control (WT) in grey on x-axis, and cell lines overexpressing BCL2 in red, or BCL2(G145E) in blue on y-axis. In case, there is no change of expression level, the data point for the GOI will reside on the diagonal within 95.0 % confidence interval shown as solid lines. A) Differential expression of transcripts with significantly increased levels of GOI upon BCL2 overexpression but opposite directionality of levels of GOI in the BCL2 mutant. pyruvate dehydrogenase kinase 3 (PDK3, Gene ID: 5165) and asparagine synthetase (ASNS, Gene ID: 440) are two highly expressed and significantly dysregulated genes identified by the screen. PDK3 is a protein kinase that regulates and deactivates the pyruvate dehydrogenase complex by phosphorylation thereby shutting off flux from lower glycolysis into the TCA cycle. ASNS supports biosynthetic and anaplerotic flux in a L-glutamine-coupled reaction, which transfers an amide from L-glutamine to aspartate to generate asparagine. B) Differential expression of transcripts with decreased levels upon BCL2 overexpression and up-regulation in the BCL2 mutant. C) Differential expression upon BCL2 overexpression but no change of directionality in the BCL2 mutant.

**Figure 4:**
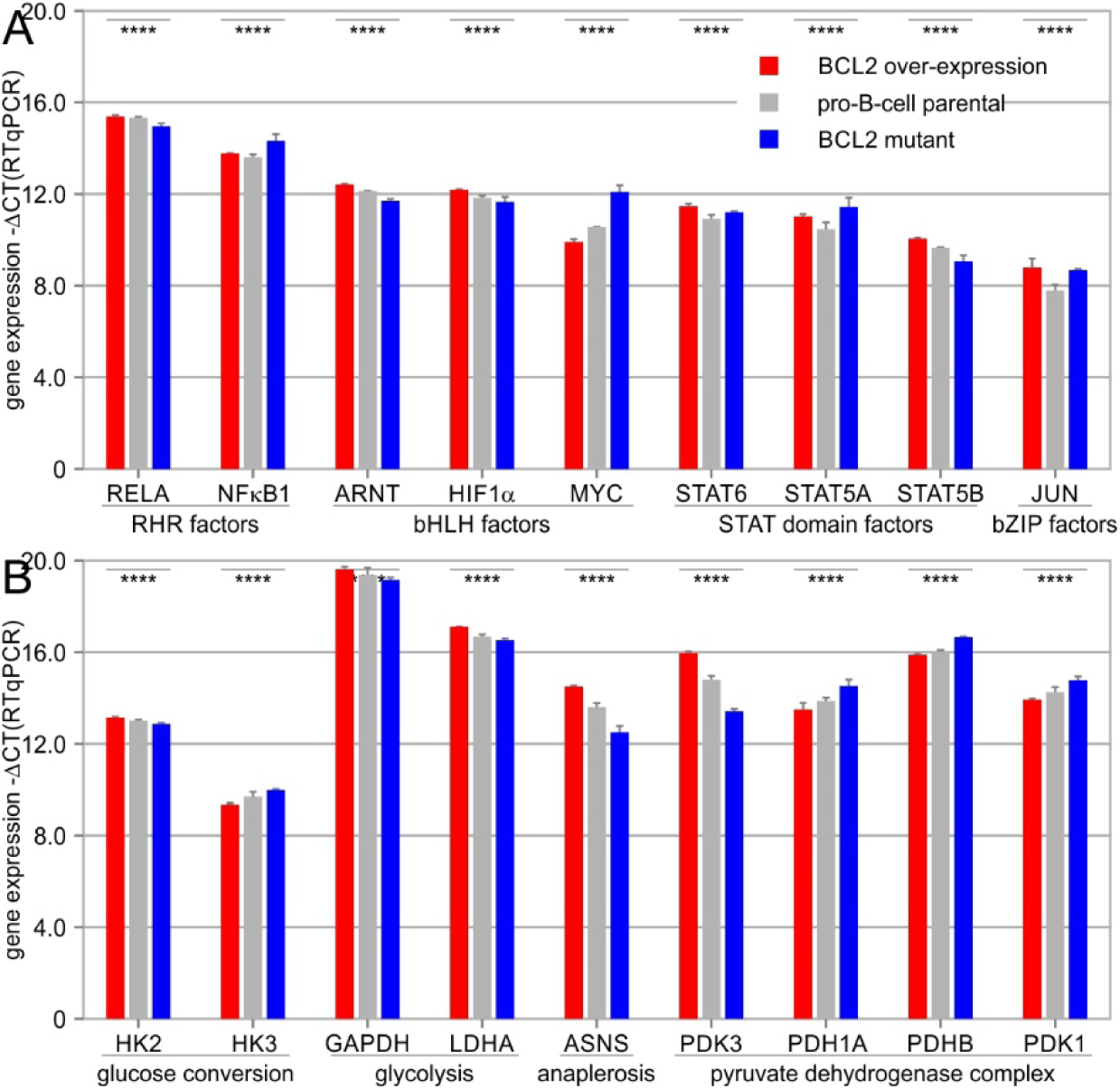
Elevated BCL2 expression causes differential expression of transcription factors and metabolic enzymes. Real-time quantitative polymerase chain reaction (RT-QPCR) validates significant deregulation of A) transcription factors and B) central carbon metabolic enzymes and their regulators. Gene expression level is shown as negative threshold cycle (-ΔCT) quantified relative to the house keeping gene with significant differential expression marked by **** and p-values below 1.0E-05.

### BCL2 overexpression increases cellular proliferation and mitochondrial mass

To characterize the effect of BCL2 overexpression on the proliferation of lymphocytic and lymphoma cell lines automated, serial cell counts were obtained. BCL2 overexpression resulted in significantly higher growth rates ρ with p-values below 1.00E-04 compared to the parental and mutant BCL2-G145E cell lines (Figure 5A). Despite an initial plateau of the overexpressing cell line, it was possible to obtain fitted exponential maximum growth rates ρ^max^ for all cell lines with an explained variation R^2^ above 0.970 of the regression model (Figure 5B). Notably, ρ can undergo dramatic changes up to 50% decrease depending on nutrient supply, cell density, and growth phase ^43^. Maximum proliferation rates at exponential growth were ρ^max^=9.59E-02 ± 2.20E-03, R2=9.73E-01, t2^max^=10.4 h for parental pro-B cell line, ρ^max^=1.08E-01 ± 3.24E-03, R2=9.83E-01, t_2_^max^=9.29 h, 112.2% increase, p-value=6.70E-03 for BCL2 overexpression, and ρ^max^=8.75E-02 ± 4.93E-04, R2=9.78E-01, t_2_^max^=1.14E+01, 10.7% decrease, p-value=5.42E-03 for the BCL2 mutant.

**Figure 5:**
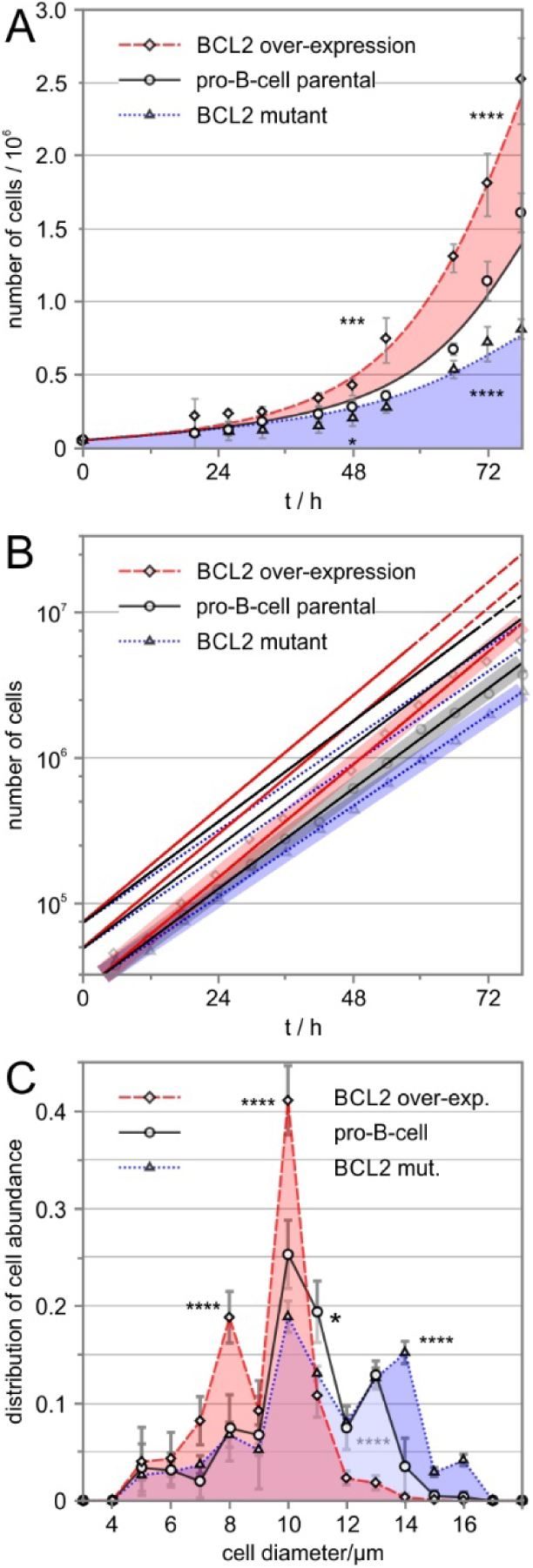
Elevated BCL2 expression significantly increases proliferation rate and decreases cell size. A) Proliferation rates of parental lymphocytic pro-B-cell lines and BCL2 overexpressing lymphoma cell lines were quantified using automated cell counting. BCL2 overexpression resulted in 12.2 % increased cellular proliferation, while expression of BCL2(G145E) mutant reduced cellular proliferation by 10.7 %. Significance levels are indicated with asterisks according to p-value thresholds (* p-value < 5.00E-02, ** p-value < 1.00E-02, *** p-value < 1.00E-03: **** p-value < 1.00E-04). B) BCL2 overexpressing lymphoma cell lines have a significantly higher fraction of cells with smaller cell diameters (8-10 μm) than parental lymphocytic pro-B-cell lines, while BCL2 mutant cells have a higher fraction of cells with larger cell diameters (14-16 μm). Distribution of cell size is shown as fraction of observed cell counts for each measured cell diameter.

Additionally, cell diameter measurements indicate that BCL2 overexpressing cells were significantly smaller on average than were parental or mutant cells. Automated cell gating with thresholds between 8-10 μm and 12-14 μm showed smaller BCL2 overexpressing cells and larger BCL2 mutant cells than the parental cell lines with p-values below 1.00E-04 (Figure 5C). Given the increased flux of D-glucose and L-glutamine into the TCA cycle, we next asked whether BCL2 overexpression affected mitochondrial dynamics more generally. Mitochondrial staining with MitoTracker green fluorescent mitochondrial stain (MTG) followed by flow cytometry revealed an increase in mitochondrial staining (Figure 6A). This is indicative of increased mitochondrial mass within BCL2 overexpressing cells, despite these cells being smaller (Figure 5C). Staining for mitochondrial membrane potential and superoxide generation by the mitochondrial superoxide indicator, MITOSOX fluorescein isothiocyanate, did not reveal any difference between BCL2-overexpressing lymphoma and the parental lymphocytic cell line (Figure 6B-C).

**Figure 6:**
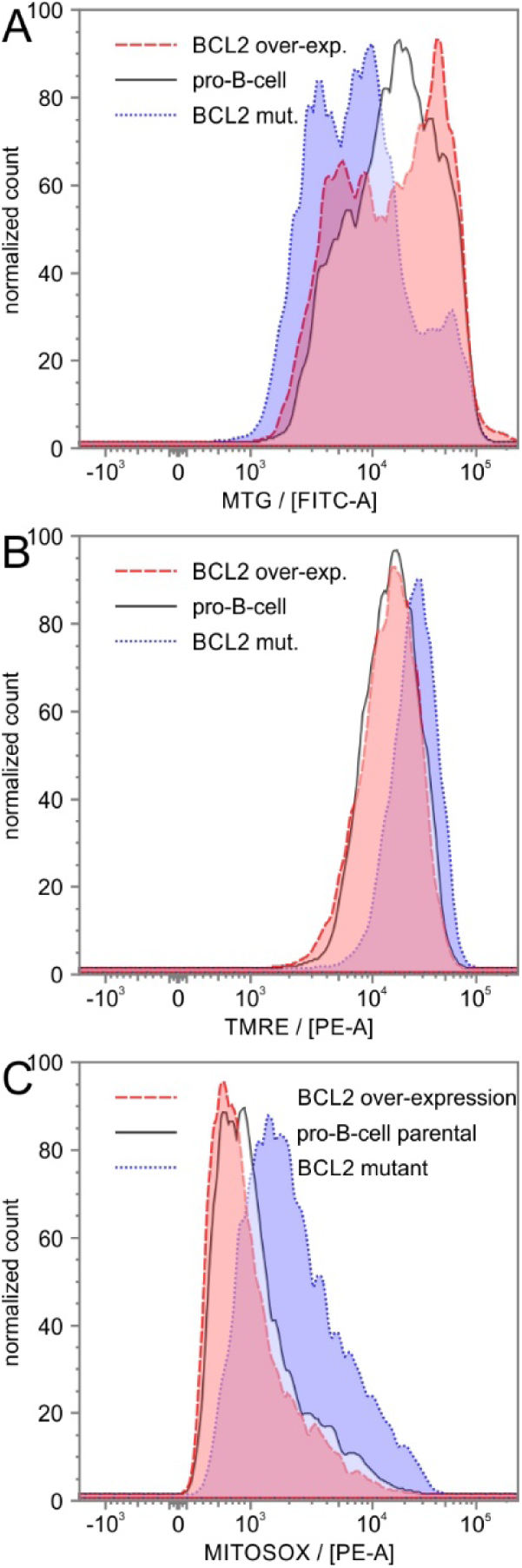
BCL2 overexpression increases mitochondrial mass. Lymphoma and parental pro-B-cell lines were treated with fluorescent mitochondrial dyes to quantify mitochondrial size, mitochondrial membrane potential, and intracellular superoxide levels by flow cytometric cell scanning. Flow cytometric cell scanning plots show normalized cell counts of 100,000 events of lymphatic and lymphoma cell lines. Markers quantified: A) fluorescence of mitochondrial tracker green (MTG, correlates with mitochondrial size, green fluorescence), B) tetramethylrhodamine ethyl ester (TMRE, determines mitochondrial membrane potential, red fluorescence), and C) mitochondrial superoxide indicator (MITOSOX, fluorescein isothiocyanate (FITC) conjugate).

### BCL2 overexpression increases glycolytic flux

Increased glycolysis is a hallmark of cancer cells and an important feature of physiological and pathologic lymphoid cell proliferation. To assess glycolytic flux in response to BCL overexpression we used GCMS-based stable isotope tracing. Following incubation with [U-^13^C_6_] D-glucose, the lymphoma cell line overexpressing BCL2 exhibited increased incorporation of labeled carbon in pyruvate and lactate compared to parental and mutant cell lines, indicating increased glycolytic and fermentative flux from D-glucose (Figure 7A-B). Moreover, BCL2 overexpression displayed increased flux of glycolytic carbon into the TCA cycle. In accordance, analysis of conditioned media samples using GCMS revealed increased D-glucose uptake and lactate secretion by the BCL2 overexpressing cell line.

**Figure 7:**
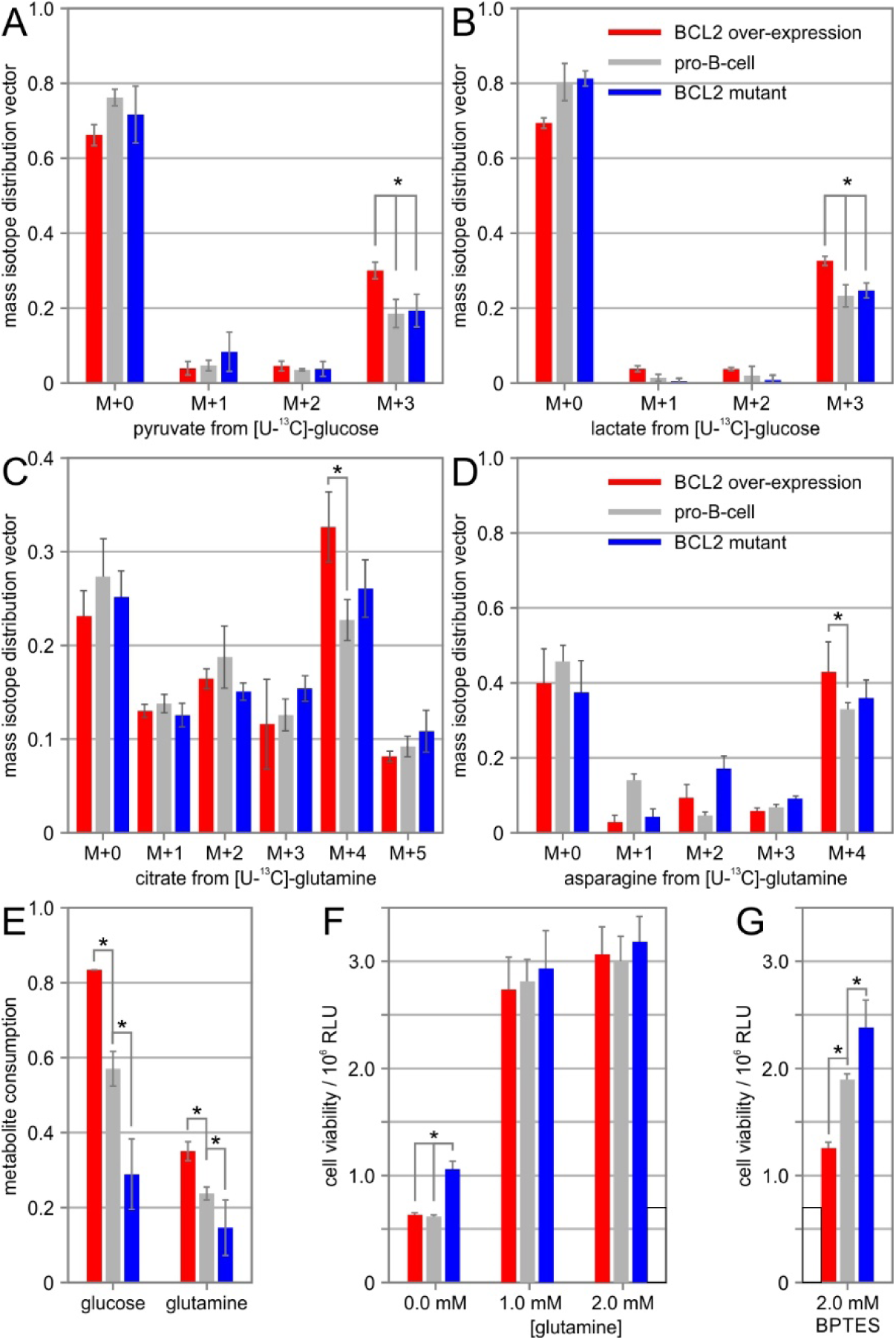
Stable isotope metabolite labeling shows increased flux into the TCA cycle and vulnerability of glutaminase inhibition. Metabolite labeling of ^13^C-D-glucose into glycolytic intermediates A) pyruvate and B) lactate of parental lymphocytic pro-B-cell lines, BCL2 overexpressing lymphoma cell lines, and cell lines with mutant BCL2 with a BH domain binding defect were quantified using gas chromatography mass spectrometry. Metabolite labeling of ^13^C-L-glutamine into TCA cycle intermediates C) citrate and D) asparagine. E) Extracellular consumption rates of D-glucose and L-glutamine. Sensitivity assay of F) L-glutamine deprivation and G) glutaminase inhibition with BPTES. Significance level of metabolic differences is indicated by asterisks based on a p-value threshold of 5.00E-02.

### BCL2 increases glutaminolysis and anaplerotic flux into TCA cycle

A prominent feature of several malignancies is a dependency on anaplerotic flux to fuel TCA and reverse TCA fluxes^35, 44^. We therefore examined whether BCL2 overexpression increased glutaminolysis and L-glutamine influx into the TCA cycle. Following incubation with [U-^13^C_5_] L-glutamine, BCL2 overexpression had increased incorporation of ^13^C stable isotopes into citrate, succinate, fumarate, L-aspartate, L-asparagine, L-glutamate and other TCA cycle intermediates compared with parental and mutant cells. In detail, the isotope composition of citrate, succinate, fumarate, aspartate, and L-asparagine showed significant increase in the M+4 feature with a p-value below 5.00E-02 in the BCL2 overexpressing cell line indicating predominantly oxidative, forward flux of the TCA cycle (Figure 7C-D). Other TCA cycle intermediates and TCA cycle associated amino acids including L-aspartate and L-asparagine track with four-carbon TCA cycle precursor organic acids characterized by reduced pool size and increased stable isotope incorporation from L-glutamine upon BCL2 overexpression. Cellular uptake of L-glutamine from extracellular media was also significantly greater in the BCL2 overexpressing cell line with p-values below 5.00E-02 (Figure 7E). Given such increased utilization of extracellular L-glutamine, we next tested whether L-glutamine deprivation preferentially affected the BCL2 overexpressing cell line. Removing L-glutamine completely from extracellular media for a period of 24 hours abolished cellular growth of pro-B-cell and BCL2 overexpressing cell lines (Figure 7F). In contrast, pro-B-cells with BCL2 mutant protein showed increased survival in the absence of L-glutamine with a p-value below 5.00E-02 (Figure 7F). Furthermore, treatment with the selective glutaminase (GLS, Gene ID: 2744) inhibitor BPTES over a period of 24 hours more adversely affected the BCL2 overexpressing lymphoma cell line in comparison to the non-tumorigenic lymphocytic cell lines with p-values below 5.00E-02 (Figure 7G). Taken together, BCL2 activation in the overexpressing cell line showed increased glycolytic and TCA cycle capacity, while creating a strong dependency on TCA cycle support via glutaminolytic anaplerosis.

## Discussion

### Pro-survival function of BCL2 beyond its anti-apoptotic role

Elevated expression of the proto-oncogene BCL2 is a key element discovered in hematological malignancies ^9, 45, 46^. The B-cell lymphocytic lineage utilizes programmed cell death as selection mechanism relying on BCL2 as deciding factor for survival or in its absence apoptosis. In response to BCL2 overexpression, pro-apoptotic BH members are bound by BCL2 preventing initiation of outer membrane permeabilization in the process of apoptotic cell death ^47^. While this observation accounts for the protective, anti-apoptotic effect of BCL2, it does not explain other aspects of BCL2 activity particularly in cancer. In malignant melanoma, myeloma, lung adenocarcinoma and stomach adenocarcinoma, overexpression of BCL2 is associated with oncogenesis and poor outcome ^49-52^. Overexpression of BCL2 in pro-B lymphocytic cell lines exhibits resistance to mitochondrial apoptosis and induces lymphoma upon injection in mice^20, 25, 48^. Correspondingly, loss of the BCL2 gene in murine models leads to widespread apoptosis, hypopigmentation, and polycystic kidney disease pointing to an important homeostatic role ^53^. Surprisingly, despite its fundamental role in apoptosis and cancer, there is little molecular or mechanistic data on the effects of BCL2 expression levels on transcriptional and metabolic networks.

### Somatic activation of BCL2 by somatic copy number amplification

In cancer, increased levels of BCL2 have been observed at the transcriptional, epigenetic, or copy number level^45, 54-56^. Congruently, the t(14:18) chromosomal translocation causes constitutive overexpression of BCL2 by juxtaposing it to immunoglobulin heavy chain gene enhancer elements. Our somatic copy number analysis identified locus chr18q21 as a frequently amplified hotspot in both, focal and arm level, genomic alterations (Figure 1, Supplementary table 1). The analysis confirmed prior described correlations between BCL2 amplification and MYC amplification, which are associated with poor prognostic outcome^43, 57, 58^. Further, we detected co-occurrence between BCL2 and NF-κB/REL signaling ^57, 59^. Curiously, frequent, focal SCNA amplification of the metabolic enzyme ASNS on chr7q21 of DLBC patients coincides with amplification of BCL2 (Figure 1A). Together, these observations raise questions for functional implications of BCL2 amplification for transcriptional and metabolic regulatory networks.

### BCL2 serves a pro-survival role by modulating and controlling genes required for apoptotic cell death

Most oncogenes, like BCL2, if they are not transcription factors themselves, are activators of transcriptional programs. Our analyses identified BCL2 overexpression as an activator of several highly regulated and inducible transcription factors, including NF-κB subunits NFKB1/2 and REL/A/B, basic helixloop-helix (BHLH) family members, HIF1A or MYC, AP1 factors, JUN and FOS, and STAT family members. Stimulation of these pathways, in turn, increases expression of target genes that are necessary for growth and protection from apoptosis. The anti-apoptotic potential of BCL2 has been demonstrated to be partially attributed to its complexing with various components of the NF-κB complex in the nucleus, thereby modulating nuclear gene expression with a strong pro-inflammatory and oncogenic outcome ^60, 61^. Since inducible transcription factors are activated by specific regulators, detection of unusually active transcription factors also points to upstream signal transduction pathways, potentially identifying therapeutic options for inhibition in BCL2-driven cancers. Our dataset contains three pieces of independent evidence—at the genomic, transcriptomic and effector target gene level—that NF-κB/REL signaling is closely connected to the oncogenic, pro-survival function of BCL2.

### The cellular model of BCL2 activation supports NFKB-positive subtypes of DLBC

In B-cell neoplasias, DLBCs represent a group of different subtypes including activated B cell-like (ABC), primary mediastinal center B cell-lymphoma (PMBL), and germinal center B cell-like (GCB) DLBC. BCL2 amplification and transcriptional activity were found in chr18q21 in ABC and PMBL ^62^. One of the most important differences among the DLBC subgroups is the constitutive activity of the NF-κB pathway in ABC and PMBL but not GCB DLBCL. Significantly, of the three DLBC subtypes, only ABC and PMBL responded to inhibitors of the NF-κB pathway, whereas GCB was impervious to these agents^62, 63^. In DLBC, JAK-STAT signaling is a feature of the ABC DLBC subtype and triggered by autocrine production of interleukins under the control of NF-κB/REL. STATs also stimulate NF-κB target genes, which could be due to the ability of STATs to form a complex with the NF-κB transcription factor complex ^64^. This emerging evidence suggests that BCL2 and NF-κB/REL positive DLBCs critically depend on sustained activity of the NF-κB pathway, which, among others, is achieved through numerous distinct genomic and transcriptomic aberrations^60, 61^. In the BCL2 overexpression model, the BCL2-NF-κB-STAT axis stood out as consistently altered at the genomic and transcriptional level emphasizing its role as important oncogenic driver pathway in lymphoma. Systems-level connections of transcriptional networks now enable us to propose target therapeutic intervention strategies that target oncogenic BCL2 signaling and dysregulated NF-κB activity.

### Pleiotropic effects of BCL2 on transcription factors

RNA-Seq enrichment and RT-QPCR validation show activation of transcription factor programs of NF-κB/REL, HIF1A/ARNT, AP1, and STAT complexes in the presence of elevated BCL2. In concordance, transcriptional profiling of cellular DLBC models, members of NF-κB, HIF, CREB and other transcription factor families were found overexpressed or inappropriately activated ^65^. The pro-B-cell lymphocytic lineage provides a useful model system independent of MYC activation. Overexpression of BCL2 in these cells was shown to suppress MYC, a target gene of NFKB1, by influencing the DNA-binding activity of NFKB1, thereby negatively affecting MYC transcription ^60^. In this case, similar to our presented model system, NFKB1 serves as upstream regulator of MYC and other transcription factors ^59^ and up-regulates MYC transcript levels in the absence of wildtype BCL2 expression (Figure 4A). Conversely, overabundance of BCL2 suppresses MYC. Independent of transcriptional regulation, there is genomic synergy between BCL2 and MYC at the copy number level (Supplementary table 1). Therefore, transcriptional control of MYC expression by BCL2 via NFKB1 can be overridden by genomic alterations of MYC, which negatively impacts patient survival. An additive adverse patient survival effect of BCL2 activation in combination with MYC or NFKB1 overexpression has been reported^66, 67^ correlating with experiments on transgenic BCL2 and MYC animal models in lymphomagenesis^68, 69^. Despite poor prognosis, MYC rearrangements are rare in BCL2-enforced lymphomagenesis in humans. Importantly, BCL2 and NFKB1 correlate at genomic, transcriptional, and network levels in DLBC patients strengthening and supporting the *in vitro* data.

### Reversal of the phenotype by BH interaction mutant of BCL2

Interactions of BCL2 with BH members regulate mitochondrial outer membrane permeabilization in the process of apoptotic cell death. In addition to canonical BH domains, structurally similar non-canonical BH domains can be found in numerous proteins, potentially making them responsive to overabundant BCL2. It is important to mention that the chosen approach of profiling in combination with enrichment analysis is not able to resolve protein-protein interactions. However, the approach prioritizes cellular networks in an unbiased way and identifies NF-κB signaling, amino acid metabolism, and inflammation as important signatures of BCL2 associated dysregulation. Further, the BCL2(G145E) mutant, which is unable to bind BH3 domains, provides a useful control, since phenotypic differences between cell lines overexpressing BCL2 wildtype vs mutant, can be attributed to an inability to bind or interact with the BH3 domain of BCL2 ^23^. Indeed, overexpression of mutant BCL2 reversed the cellular phenotype observed with wildtype BCL2 overexpression. This indicates that lack of BH domain interaction in the mutant reduces proliferation rate, increases cell size, and results in smaller mitochondria compared with wildtype BCL2 or control pro-B cell lines (Figure 5-6). Similarly, transcriptomic and metabolic assays point to phenotypic nodes, which are responsive to BCL2 overexpression and which can be reversed by the interaction mutant. Specifically, transcription factors RELA, ARNT, HIF1A, and STAT5B correlate positively with an increase in BCL2 expression and BH interactions (Figure 3A and 4A). Similarly, transcriptional levels of metabolic effector enzymes HK2, GAPDH, LDHA, ASNS, and PDK3 follow BCL2 activity (Figure 3A and Figure 4B).

### Dysregulated amino acid and TCA cycle metabolism as part of the BCL2 effector network in cancer

The integral relationship between BCL2 and mitochondria suggests that it may function in regulating cellular metabolism independent of its role in apoptosis. Stable isotope-assisted metabolic flux measurements showed that elevated BCL2 expression increases carbon utilization necessary to support cell cycle progression and cellular proliferation. Tumorigenic overexpression of BCL2 significantly increased flux from D-glucose into pyruvate and lactate, which is indicative of oxidative fermentation and the Warburg effect in cancer cells. Transcriptional networks responding to BCL2 overexpression include activation of NF-κB, HIF1A, STAT5A and AP1 target genes ensuring activation of fermentative glycolysis and oxidative glutaminolysis (Table 1). The glycolytic phenotype is carried out by transcriptional up-regulation of rate-limiting enzymes HK2, GAPDH, and LDHA. The pentose phosphate pathway is increased, judged by up-regulation of gate-keeper enzymes G6PD, TKT, and TALDO1. The transcription factor complex HIF1A/ARNT controls glycolytic enzyme levels and supports rapid growth. However, by deregulating PDK3, which is another HIF1A target gene, the mitochondrial acetyl-CoA pool is separated from the glycolytic one. The TCA gatekeeping pyruvate dehydrogenase (PDH) complex is switched off by elevated activity of PDK3, which causes inhibition of the PDH complex. The activator of PDHs, PDP2, as well as enzymatic subunits of PDH, PDHA1, and PDHB, are down-regulated to prevent the irreversible conversion from pyruvate into acetyl CoA. At the same time, any other PDK isoforms, PDK1, PDK2, and PDK4, are down-regulated emphasizing PDK3 as sole regulatory step for switching off PDHs. Decoupling of glycolysis depletes TCA cycle metabolite pool sizes and increases dependence on anaplerosis to replenish TCA cycle intermediates ^44^. Anaplerotic L-glutamine metabolism is engaged via up-regulation of GLS and ASNS to compensate and support mitochondrial TCA cycle metabolism. The tight connection between BCL2 and ASNS is illustrated by co-occurring SCNAs, synergistic transcriptional regulation, joint strong up-regulation at the protein level, and increased metabolic flux (Figure 1A, 2B, 3A, 4B, 7D). ASNS supports biosynthetic and anaplerotic flux in an L-glutamine-coupled reaction by transferring an amide group from L-glutamine to aspartate thereby generating L-asparagine. L-glutamine, α-ketoglutarate, and L-asparagine metabolism is not only important for cancer cell survival but also in oxidative stress and tumor vascularization to offset nutrient and oxygen limitations.

### Metabolic bottlenecks upon BCL2 overexpression and dependency on anaplerotic support

Combined metabolic and transcriptional analyses highlight distinct switches attributed to overexpression of BCL2. Enhanced BCL2 expression and BH binding scaffolds stimulate pro-survival interactions and promote mitochondrial stability. In addition, BCL2 overabundance stimulates non-canonical nuclear interactions leading to activation of transcriptional networks facilitating onco-metabolism. Of the tested transcription factors, transcriptional complexes of NFKB1, RELA, HIF1A, ARNT, and STATs are prominently expressed and able to modulate expression of metabolic target genes. Beyond the traditional Warburg effect, which is focused on cytosolic fermentation, the TCA cycle-gatekeeper, PDK3, separates glycolysis and mitochondrial TCA cycle. Congruently, anaplerotic L-glutamine and L-asparagine conversion supports oxidative TCA cycle metabolism and fuels elevated bioenergetics demands of rapidly growing BCL2-positive cells. Since the discovery of L-asparagine-sensitive lymphomas ^70^, a great deal of research has probed the nature of L-asparagine turnover in cancer and its value as a therapeutic target ^71^. In BCL2 overexpressing cell lines, targeting of L-glutamine turnover was immediately responsive, likely due to a metabolic bottleneck created by increased dependency on anaplerotic support. Moreover, ^13^C-assisted stable isotope tracing showed how BCL2-driven lymphoma cell lines are able to reroute TCA cycle related amino acid metabolism and sustain L-glutamine and L-asparagine pathways. Details of how anti-apoptotic regulation generates specific conditions under which non-essential amino acids becomes indispensable for cancer cells deserve future attention.

## Conclusion

The field of BH domain interactions is at an exciting point. Extensive knowledge and datasets including high-resolution structural and systems biology allow to build on decades of research on pro- and anti-apoptotic mediators providing useful insights into transcriptome, proteome, and interactome of the BH family. Importantly, the therapeutic potential of BCL2 activation by somatic amplification or transcriptional up-regulation has been recognized and offers a genotype-match approach for precision targeting of cancers affected by dysregulation in the pro-survival BH family of onco-proteins. These results suggest DLBC subtype-specific biomarkers based on genomic and transcriptomic alterations of BCL2 and encourage stratification of DLBC patients for targeted therapy with BCL2 inhibitors. Based on genomic, transcriptional, and metabolic readouts of BCL2-activated lymphoma, BCL2 has a pronounced oncogenic phenotype. By triggering a non-canonical network of transcription factors it promotes a metabolic and mitogenic program. Anaplerotic L-glutamine metabolism is engaged via up-regulation of GLS and ASNS creating metabolic vulnerability. Significantly, there is an increased sensitivity to glutaminase inhibition and glutamine deprivation in BCL2-driven lymphoma cells. This is supported by ^13^C-assisted metabolomics data indicating increased anaplerotic glutamine and asparagine flux. The NF-κB/REL complex stands out as master regulator of pro-inflammatory and mitogenic target networks. In addition, it is controlling and impacting downstream transcriptional networks of stimulated STAT and repressed MYC transcription factor networks. The BCL2-NF-κB-STAT axis recommends itself as biomarker and anti-cancer target to address an unmet clinical need for the precision management of lymphoma. Dual, BCL2- and NF-κB-expressing lymphomas identify a distinct molecular subset of DLBC. In the treatment of such dual BCL2- and NF-κB-expressing lymphomas, agents targeting BH domain interactions, immunotherapy, and chemotherapy focused on metabolic vulnerability should offer significant therapeutic benefit.

## Declarations

### Ethics approval and consent to participate

All experimental protocols were approved by the Institutional Review Boards at the University of California Merced. The study was carried out as part of IRB UCM13-0025 of the University of California Merced and as part of dbGap ID 5094 for study accession phs000177.v9.p8 on somatic mutations in cancer and conducted in accordance with the Helsinki Declaration of 1975.

